# A novel antiviral formulation inhibits a range of enveloped viruses

**DOI:** 10.1101/2020.03.29.009464

**Authors:** Nicola F. Fletcher, Luke W. Meredith, Emma Tidswell, Steven R Bryden, Daniel Gonçalves-Carneiro, Yasmin Chaudhry, Claire Shannon Lowe, Michael A. Folan, Daniella A Lefteri, Marieke Pingen, Dalan Bailey, Clive S. McKimmie, Alan W. Baird

## Abstract

Some free fatty acids derived from milk and vegetable oils are known to have potent antiviral and antibacterial properties. However, therapeutic applications of short to medium chain fatty acids are limited by physical characteristics such as immiscibility in aqueous solutions. We evaluated a novel proprietary formulation based on an emulsion of short chain caprylic acid, ViroSAL, for its ability to inhibit a range of viral infections *in vitro* and *in vivo. In vitro*, ViroSAL inhibited the enveloped viruses Epstein-Barr, measles, herpes simplex, Zika and orf parapoxvirus, together with Ebola, Lassa, vesicular stomatitis and SARS-CoV-1 pseudoviruses, in a concentration- and time-dependent manner. Evaluation of the components of ViroSAL revealed that caprylic acid was the main antiviral component; however, the ViroSAL formulation significantly inhibited viral entry compared with caprylic acid alone. *In vivo*, ViroSAL significantly inhibited Zika and Semliki Forest Virus replication in mice following the inoculation of these viruses into mosquito bite sites. In agreement with studies investigating other free fatty acids, ViroSAL had no effect on norovirus, a non-enveloped virus, indicating that its mechanism of action may be via surfactant disruption of the viral envelope. We have identified a novel antiviral formulation that is of great interest for prevention and/or treatment of a broad range of enveloped viruses.

## 1.1 Introduction

The antimicrobial properties of fatty acids have been extensively reported in the literature (for review, see Thormar et al. (Thormar and Hilmarsson, 2007) and (Churchward et al., 2018). Previously, (Thormar et al., 1987) demonstrated the antiviral effects of 14 different free fatty acids and lipid extracts from human milk against vesicular stomatitis virus (VSV), herpes simplex virus (HSV) and visna virus revealed that short chain saturated fatty acids (butyric, caproic and caprylic) together with long chain saturated fatty acids (palmitic and stearic) had no or very little antiviral activity, whereas medium chain saturated entities including capric, lauric, myristic and long chain unsaturated oleic, linoleic and linolenic acids were anti-viral, albeit at different concentrations. Another study (Hilmarsson et al., 2005) reported similar trends in the antiviral activity of six medium chain fatty acids together with their alcohol and mono-glyceride derivatives against herpes simplex viruses 1 and 2. In contrast, Dichtelmuller et al (2002) reported that caprylic acid had antiviral activity against enveloped viruses including human immunodeficiency virus, bovine viral diarrhoea virus, Sindbis virus and pseudorabies virus (Dichtelmuller et al., 2002, Pingen et al., 2016). Studies investigating the antiviral properties of whole milk reported no antiviral properties of fresh human milk, whereas milk that had been stored at 4°C possessed potent antiviral activity against several viruses *in vitro*. Refrigeration disrupts the milk fat globule membrane allowing ingress of milk serum lipase which results in hydrolysis of milk fat triglyceride (Thormar et al., 1987, Isaacs et al., 1990). It was concluded that release of fatty acids from milk triglycerides in stored milk, and that recovered from neonatal (achlorhydric) stomachs, was responsible for generating antiviral factor(s) (Thormar et al., 1987).

We investigated the effect of a specifically formulated emulsion of free fatty acids, ViroSAL, on infectivity of enveloped and non-enveloped viruses. Caprylic acid delivered in the ViroSAL emulsion exhibited significant anti-viral effects. A range of enveloped viral infection systems was utilized, and complete inhibition of viral infection was observed without any evidence of cytotoxicity. ViroSAL had no effect on the infectivity of a non-enveloped virus, norovirus, which is in agreement with previous studies demonstrating that free fatty acids are ineffective against non-enveloped viruses (Thormar et al., 1987, Kohn et al., 1980). Furthermore, ViroSAL inactivated the enveloped mosquito-borne viruses Semliki Forest virus (SFV) and Zika virus (ZIKV) *in vitro*. Prophylactic topical treatment of viral infection in mosquito bites with ViroSAL inhibited local replication and dissemination of SFV and plasma levels of ZIKV in mice. Transmission electron microscopy analysis indicated that ViroSAL disrupts orf parapoxvirus envelope integrity, with higher concentrations completely disrupting virion morphology. These data indicate that ViroSAL has antiviral activity against a range of enveloped viruses in vitro and in vivo.

## 1.2 Materials and Methods

### 1.2.1 Cell lines and antibodies

HepG2, Caco-2, A549, Vero cells, RAW 264.7 murine macrophages, 293T human embryonic kidney cells and BV2 murine microglial cells were propagated in DMEM supplemented with 10% fetal bovine serum (FBS), 2 mM L-glutamine (Gibco), antibiotics (100U of penicillin/mL and 100μg of streptomycin/mL) and 1% non-essential amino acids (Gibco). Baby hamster kidney– 21 (BHK-21) cells and C6/36 mosquito cells were grown as previously described (Pingen et al., 2016). Vero cells engineered to over-express signaling lymphocyte activation molecule (SLAM) were supplemented with 0.4mg/mL of geneticin (G418, Sigma). Human foreskin fibroblasts (HFF-1) cells were obtained from ATCC (ATCC^®^ SCRC-1041™), and were propagated in DMEM supplemented with 15% FBS, 2mM l-glutamine, antibiotics (100U of penicillin/mL and 100μg of streptomycin/mL) and 1% non-essential amino acids). Primary tonsil epithelial cells were isolated from normal tonsil tissue obtained during hypophysectomy, as previously described (Feederle et al., 2007). Primary foetal lamb skin (FLS) cells were isolated as previously described (Scagliarini et al., 2007) and cultured in Medium 199 with 10% heat inactivated FBS. All cells were maintained at 37°C, 5% CO_2._

Anti-HSV-1 gD antibody was a generous gift from Colin Crump, University of Cambridge (Minson et al., 1986). Anti-mouse alexa-488 antibody was obtained from Gibco, U.K.

### 1.2.2 Pseudovirus generation and in vitro infection

Pseudoviruses were generated by transfecting 293T cells with plasmids encoding a HIV-1 provirus expressing luciferase (pNL4-3-Luc-R-E-) and vesicular stomatitis virus (VSV-G), Zika, Ebola, Lassa or SARS coronavirus-1 (SARS-CoV-1) envelope or a no envelope control (Env-), as previously described (Fletcher et al., 2015(Broer et al., 2006). Zika virus pseudoviruses were generated by amplifying preM-M-Env region (positions 753-3549) from the PE243 Brazil Zika strain (Accession No. KX197192), with flanking restriction sites 5’NheI-insert-XbaI-3’ then inserting into pcDNA3.1(+) plasmid.

Virus containing media was incubated with the indicated concentrations of ViroSAL, at pH5.5, for 2 minutes at room temperature unless stated otherwise, and an equal concentration of neutralizing buffer was added to restore pH to 7, and virus/ViroSAL solutions were added to target cells for 8 hours. Each preparation of ViroSAL was titrated in the relevant media used to culture each cell type (DMEM, Williams E and M199). To control for pH treatment, virus was treated with pH5.5 buffer that did not contain ViroSAL for 2 minutes and then pH restored to 7 as described for ViroSAL treatments and virus added to target cells. After 8 hours, unbound virus was removed and the media replaced with DMEM/3% FBS. At 72h post-infection the cells were lysed, luciferase substrate added (Promega, U.K.) and luciferase activity measured for 10 seconds in a luminometer (Lumat LB9507). Specific infectivity was calculated by subtracting the mean Env-pp RLU signal from the pseudovirus RLU signals. Infectivity is presented relative to untreated control cells by defining the mean luciferase value of the replicate untreated cells as 100%. To assess cell viability following ViroSAL treatment, an MTT assay was performed on replicate wells in every experiment (Gibco, U.K.).

### 1.2.3. Virus culture and infection assays

Recombinant measles virus (MeV) strain IC323 carrying the open reading frame of the enhanced green fluorescent protein (EGFP) was generated as described previously (Hashimoto et al., 2002). After initial recovery, MeV was produced in Vero-SLAM cells using an initial multiplicity of infection of 0.1. When infection was fully developed, flasks were vigorously shaken and supernatants collected and clarified at 3500rpm for 30min at 4°C. Supernatant was collected, aliquoted and stored at −80°C.

Recombinant wild-type Epstein-Barr virus (EBV 2089) with a GFP insert (Delecluse et al., 1998) was generated from 293T cells carrying the recombinant EBV B95.8 genome, as previously described (Shannon-Lowe et al., 2006). Encapsidated and enveloped virus was purified from the culture supernatants by centrifugation on an OptiPrep (Axis Shield) self-generated gradient (Shannon-Lowe et al., 2009) and quantitated by quantitative PCR (qPCR) of a single-copy gene, *BALF5*, as previously described (Shannon-Lowe et al., 2006).

Herpes simplex virus (HSV-1)(Strain I7) was propagated in Vero cells as previously described (Ren et al., 2012). Titration was performed on confluent cultures of Vero cells with a 50% agarose overlay in DMEM/2% FCS, and infectious titre determined in plaque forming units (PFU).

For *in vitro* work, Zika virus (Brazil, PE243) was propagated in Vero cells, as previously described (Chavali et al., 2017). Titration was performed on confluent cultures of Vero cells with a 50% agarose overlay in DMEM/2% FCS and infectious titre quantified in plaque forming units (PFU). Virus stocks were pooled and titred on Vero cells to determine IC50 values.

Orf virus (MRI-scab) was propagated by scarification of lambs as previously described (McInnes et al., 2001) and then cultured by infection of primary foetal lamb skin (FLS) cells. Infection experiments were performed by direct inoculation of orf virus onto primary cultures of FLS cells (Scagliarini et al., 2007).

Murine norovirus (MNV-1.CW1 strain) was propagated in murine RAW 264.7 cells (Wobus et al., 2004). The yield of infectious virus was determined at 24h post-transfection of cDNA or capped RNA. Titres of virus were determined as 50% tissue culture infectious dose (TCID50) in RAW 264.7 cells, using microscopic visualization for the appearance of cytopathic effect.

The pCMV-SFV4 and pCMV-SFV6 backbone for production of SFV has been previously described (Ulper et al., 2008). SFV plasmids were electroporated into BHK-21 cells to generate infectious virus. SFV4 is the prototypic, less virulent strain of the virus, whereas SFV6 is a copy of a virulent strain (Ferguson et al., 2015).

For all *in vivo* experiments with SFV and ZIKV (Brazil, PE243), viruses were grown once in BHK-21 cells and then passaged once in C6/36 *Aedes* mosquito cells and titrated before use because mosquito cell–derived virus has distinct glycosylation and because insect cells impose distinct evolutionary constraints on viral progeny (Moser et al., 2018). SFV4 and SFV6 were used at passage 2.

### 1.2.4 Construction of ViroSAL

ViroSAL was a gift for research purposes from Westgate Biomedical Ltd, Donegal, Ireland, (Folan M. Patent WO 2011/061237). All components used in the construction of ViroSAL are pharmaceutical grade constituents of greater than 97% purity. The ViroSAL emulsion used in these studies was constructed using Lipoid S75 Lecithin (Lipoid AG, Steinhausen, Switzerland) 1.26 % W/W which had been de-lipidised using solvent extraction to remove any extraneous lipid that remained conjugated to its lipophilic sites (de-lipidised lecithin is amphipathic). A co-surfactant Pluronic F-68 (BASF Cork, Ireland) was used to enhance stability and a di-palmitoyl, 1,2-Dipalmitoyl-*sn*-glycero-3-phosphorylglycerol sodium salt (DPPG from Lipoid AG) was used to amplify surface charge on the emulsion droplet. Pharmaceutical grade caprylic acid (Merck, Nottingham, UK) 10% W/W was emulsified in the de-lipidised lecithin, Pluronic and DPPG mix 0.45% W/W and 0.54% W/W respectively using an Emulsiflex C-5 (Avestin, Ottowa, Canada) at 1,000 Bar pressure. The emulsion was diluted to 2% W/W (0.2% caprylic acid), or used at the concentrations specified in each experiment, in sodium citrate buffer, pH 5.0, with a physiological isotonicity of 280mOsM, optimized for each culture medium used, and autoclaved before use (Laverty et al., 2015).

For topical application to mouse skin, a gel formulation was constructed using 1%W/V Carbopol 974p (Lubrizol Inc, Cleveland, Ohio, USA), dispersed and hydrated in an aqueous solution of 15% glycerol (Merck) which was then pH adjusted to 5.0 before addition of 10% ViroSAL emulsion. The gel was applied liberally to test sites as described.

### 1.2.5 Assessment of the effect of ViroSAL on viral infection *in vitro*

ViroSAL has an acidic pH which necessitated minimal time exposure to cell lines followed by a neutralizing buffer to restore pH to 7.0. In this study, ViroSAL at the indicated concentrations was mixed with an equal volume of viral inoculum (MeV:original TCID50 = 4.48^7^/mL, HSV-1: 100 PFU/mL, EBV: MOI=10, Zika: MOI=10, Orf: 4 PFU/mL) or pseudoviruses bearing VSV, Ebola, Lassa or SARS-CoV-1 envelope glycoproteins, and incubated at room temperature for 2 minutes. The same volume of a buffered neutralizing solution was added and mixed thoroughly to restore the pH to 7. ViroSAL and neutralizing solution was optimized for each culture medium used in *in vitro* studies. As a control for the effect of low pH on infectivity, 200µL of pH5.5 buffer solution was added to the virus, incubated for 2min and neutralized as before. Virus/ViroSAL or control treated virus was inoculated onto appropriate target cells and incubated for 48h at 37°C, then fixed and infection enumerated, or, for pseudovirus assays, lysed and luciferase activity quantified as previously described (Fletcher et al., 2015). For MeV, TCID50s were calculated in VeroSLAM cells in triplicate using the Reed-Muench method (Reed, 1938).

For non-enveloped viral infectivity assays, a similar approach was used to enveloped virus assays. ViroSAL was mixed with murine norovirus (MNV-CW1) in a 1:1 ratio, incubated at room temperature for 2 minutes and the pH was restored to 7. Virus infection of permissive RAW 264.7 (mouse macrophage) and BV2 (murine microglial) cells were conducted immediately after neutralization in triplicate, incubated at 37°C and quantified 96h post infection using TCID50 via microscopic visualization for the appearance of cytopathic effect.

### 1.2.6 Assessment of the effect of ViroSAL on viral infection in vivo

6-8-week-old female C57bl/6J mice were derived from a locally bred colony maintained in a pathogen-free facility, in filter-topped cages, and maintained in accordance with local and governmental regulations. To prevent genetic drift, mice have been rederived using externally supplied mice (Charles River).

Because arbovirus infection of the skin always occurs in the context of an arthropod bite, we used a mouse model that additionally incorporates biting *Aedes aegypti* mosquitoes. Host response to mosquito bites includes oedema and an influx of leukocytes that enhances host susceptibility to infection with virus (Pingen et al., 2016). Therefore, we used our previously established model of arbovirus infection at mosquito bites. This model was specifically developed to model natural infection by arbovirus, including mimicking the same dose delivered by mosquitoes, using mosquito cell–derived virus, injecting a small 1ul inoculum volume, and by including the presence of a mosquito bite at the site of inoculation. To ensure that mosquitoes bit a defined area of skin (upper side of the left foot), anesthetized mice were placed for up to 10 min into a mosquito cage containing *A. aegypti* mosquitoes (locally bred colony derived from the Liverpool strain). Biting was restricted to a defined area of the left foot by covering all other mouse skin with an impenetrable barrier.

Following mosquito biting, a preparation containing 10%W/W ViroSAL was applied to the mosquito bite. 20 minutes post ViroSAL application, the *A. aegypti* bites were inoculated with 1ul 0.75% BSA PBS solution containing 2×10^4^ SFV4 by hyperfine needle (Hamilton). ViroSAL was re-applied to the inoculation site 5 hours post-infection. 24 hours post infection, mice were culled and blood, spleen and the skin of the inoculation site harvested. For Zika virus infections, to allow successful viral infection, mice were administered 1.5mg of IFNAR mAB (InVivoMAb anti-mouse IFNAR-1 Clone: MAR1-5A3) subcutaneously in the upper back, 24 hours prior to infection. Note, infected mosquitoes were not used to inoculate virus to mice because the inoculum supplied by biting mosquitoes is too variable and unpredictable to allow effective comparisons.

Viral RNA and host gene transcripts in tissues were quantified by reverse transcription qPCR, and infectious virus was quantified by end point titration, as described previously. Tissue generated up to 100 ug of total RNA, of which 1 ug of RNA was used to create cDNA, of which 1% was used per qPCR assay. qPCR primers for SFV amplified a section of E1 and primers for ZIKV amplified a section of the env gene. For SFV and ZIKV, qPCR assays measured the sum value of both genome and subgenomic RNA (Bryden et al., 2020). Each result represents the median of three or four technical replicates of one biological replicate. For plaque assay, viral stocks and biological samples were serially diluted, and each dilution was assayed in duplicate on BHK-21 cells with an Avicell overlay for 48 hours as previously described (Pingen et al., 2016). Biological replicates from mice were excluded from analysis if injection of virus inadvertently punctured a blood vessel (although this was rare and occurred with a frequency of <1%).

All experiments involving mice had been subject to rigorous review by the University of Leeds welfare and ethical review committee and additionally approved by the U.K. Home Office (license PA7CF4E75).

### 1.2.7 Flow Cytometry

Zika infected cells were fixed with 4% paraformaldehyde, stained with primary anti-Zika-E-protein antibody (1:1000, 4G2), and secondary anti-mouse Alexa-488 (1:1000). Cells were then resuspended in PBS, and 10,000 cells were screened by flow cytometry using a FACScalibur Flow Cytometer (BD Biosciences, Germany). Data was analyzed using FlowJo Software (FlowJo LLC, USA).

### 1.2.8 Transmission electron microscopy

Orf (MRI-scab) was harvested from supernatants of primary cultures of foetal lamb skin (FLS) cells, filtered through a 0.45μm filter and then centrifuged for 12 hours at 12,000rpm in a bench-top centrifuge (Eppendorf). Virus was placed on formvar coated copper 400 electron microscopy grids for 5 minutes; excess supernatant was removed and the grids were stained with 1% uranyl acetate. Grids were visualized using a Tecnai 12 transmission electron microscope.

### 1.2.9 Statistical analysis

*In vitro* results are expressed as the mean ± 1 standard deviation of the mean (SD), except where stated. Statistical analyses were performed using Student’s t-test or Mann-Whitney U test in Prism 4.0 (GraphPad, San Diego, CA) with a P <0.05 being considered statistically significant. For all in vivo-derived data, data were analyzed using GraphPad Prism Version 7 software. Copy numbers of viral RNA and infectious titres from virus-infected mice were not normally distributed (with data points often spread over orders of magnitude) and were accordingly analyzed using the nonparametric-based Mann-Whitney test or Kolmogorov-Smirnov test.

## 1.3 Results

### 1.3.1 ViroSAL inhibits a diverse range of pseudoviruses

To assess the ability of ViroSAL to inhibit viral infection, we utilized a pseudovirus system, which allows high throughput assays to assess viral entry to cells. ViroSAL inhibited entry of viral pseudoparticles bearing the envelope glycoproteins of VSV, Lassa, Ebola and SARS-CoV-1 viruses. Pseudoviral particles mirror the entry pathways of their respective wild type viruses and use the same viral receptors and entry pathways (for review, see (Li et al., 2018)). These constructs generate high titre pseudoviruses that can be used to infect a range of cell types. Inhibition of pseudovirus entry by ViroSAL occurred in a concentration-dependent manner, when pseudoviruses were incubated with ViroSAL for 2 minutes (Figure 1A). Treatment of pseudoviruses with 2% ViroSAL resulted in complete neutralization of pseudovirus infectivity on 293T human embryonic kidney cells (Figure 1A). There was no alteration in cellular proliferation or cytotoxicity when cells were treated with pseudovirus or ViroSAL, assessed using an MTT assay (data not shown). ViroSAL is stable at pH5.5, so pseudovirus treatment was carried out at pH5.5 and then restored to pH7 before infection of eukaryotic cells. Treatment of VSV and Lassa pseudoviruses with pH5.5 control buffer caused no significant change in infectivity; however, a significant reduction in Ebola pseudovirus infectivity was observed (Figure 1A).

**Figure 1:**
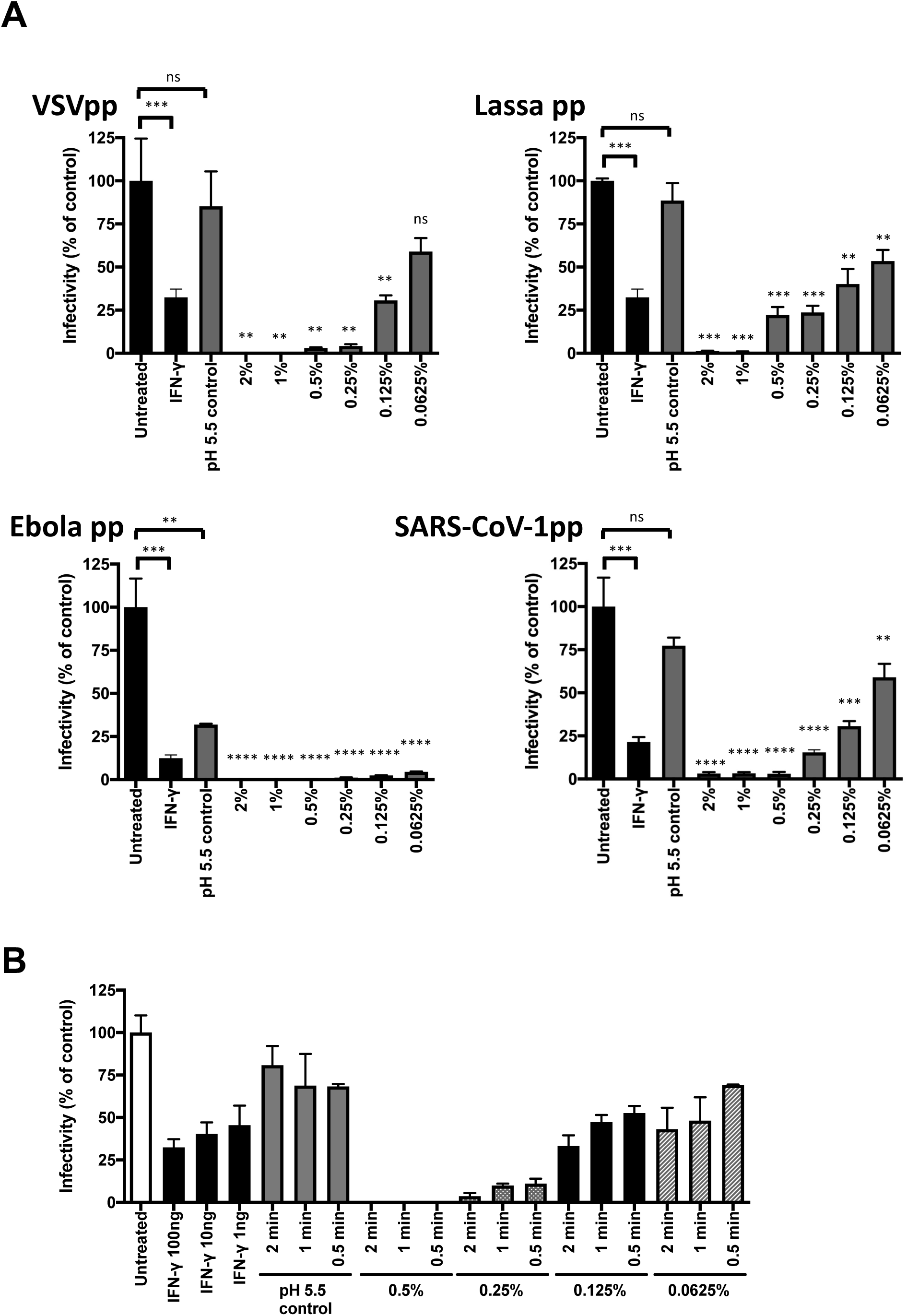
ViroSAL inhibits cellular entry of a diverse range of enveloped pseudoviruses. (A) Pseudovirus bearing the envelope glycoproteins of VSV, Lassa, Ebola or the spike protein of SARS-CoV-1 was treated in a 1:1 dilution with concentrations of ViroSAL ranging from 4% to 0.125% (final concentrations ranged from 2% to 0.0625%) for 2 minutes. Buffer was then added to restore the pH to 7. To control for the effect of pH on viral infectivity, virus was treated with a pH5.5 buffer, equivalent to that of ViroSAL, for 2 minutes and then the pH of the virus was restored to 7. Pseudoviruses were used to infect 293T human embryonic kidney cells. 10ng/ml IFN-y was included as a positive antiviral control. Data are presented as mean infectivity ± SD relative to the untreated virus control. (B) Pseudovirus bearing the envelope glycoproteins of VSV were treated for 2, 1 or 0.5 minutes with ViroSAL, the pH restored to 7, and used to infect 293T cells. Control virus was treated with a pH5.5 buffer for 2 minutes and the pH restored to 7. Statistical differences are presented relative to the pH5.5 control (n = 3 independent experiments). \*\*\*\**P* < 0.0001, \*\*\**P* < 0.001, \*\**P* < 0.01, ns: not significant.

To establish whether ViroSAL inhibits pseudovirus infection in other cell types, ViroSAL-treated Lassa and VSV pseudoparticles were used to infect human intestinal (Caco-2) and hepatoma (HepG2) epithelial cell monolayers. Similar levels of neutralization with ViroSAL were observed compared with that of 293T cells. The inhibition of viral infectivity with pH5.5 control treatment was perhaps due to the less efficient entry of viral pseudoparticles to these cells compared with the highly permissive 293T cell line (Supplementary Figure 1). Because many aspects of enveloped virus lifecycles are pH sensitive (Ruigrok et al., 1992) and pH is the most common variable when assessing the antimicrobial activity of fatty acids (Churchward et al., 2018), in all cases the control against which ViroSAL treatments were compared were those exposed to virus at pH 5.5.

### 1.3.2 ViroSAL inhibits pseudovirus infection in a time-dependent manner

To test whether ViroSAL inhibits pseudovirus infection following short treatment times, VSV pseudoviruses were incubated with ViroSAL at concentrations from 0.0625% to 0.5% for 2, 1 or 0.5 minutes, and then used to infect 293T cells. A time-dependent decrease in infectivity was observed (Figure 1B), and ViroSAL inhibited VSV pseudovirus at concentrations greater than 0.25% following 30 seconds of treatment.

### 1.3.3 ViroSAL efficiently inhibits a range of infectious enveloped viruses

To test whether ViroSAL is capable of inhibiting full-length infectious viruses, we used a range of infectious virus systems and selected cell culture systems. Pseudoviruses were generated bearing the envelope glycoproteins of Zika virus strain PE243, a Brazilian strain, and the ability of ViroSAL to inhibit pseudovirus entry together with the full-length infectious clone of Zika virus strains PE243 and MR766, an African strain, were compared. ViroSAL inhibited both Zika virus pseudovirus (Figure 2A) and infectious virus (Figure 2B,C), although a higher concentration of ViroSAL (3%) was required for complete neutralization of full-length viruses compared to pseudoviruses.

**Figure 2:**
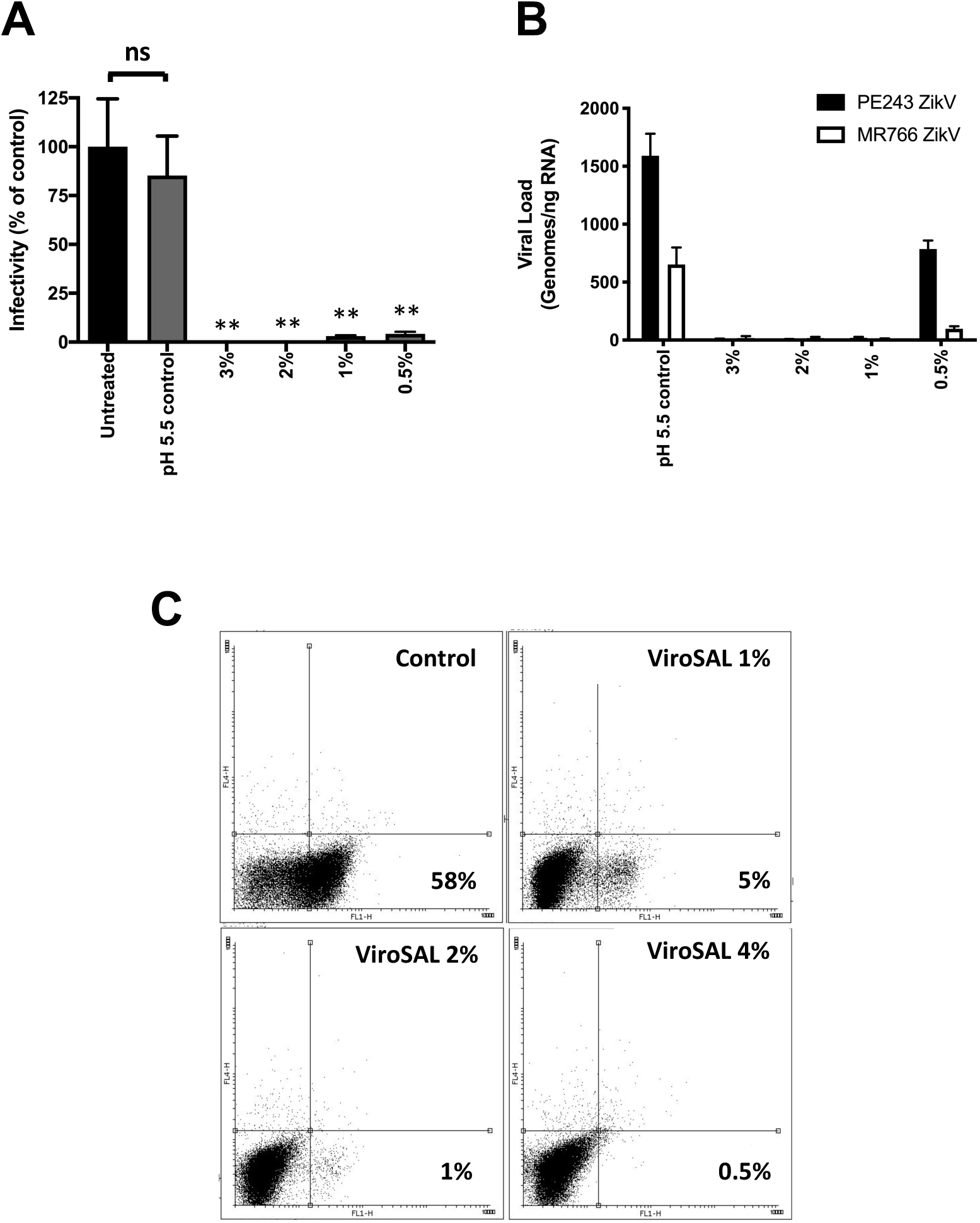
ViroSAL inhibits Zika pseudovirus and full-length infectious virus. (A) Zika pseudovirus was treated with a range of dilutions of VIROSAL for 2 minutes, the pH restored to 7, and the pseudoviruses were added to human foreskin cells. Control virus was treated with a pH5.5 buffer for 2 minutes and the pH restored to 7. (B) Two infectious clones of Zika virus, PE243 (Brazilian strain) and MR766 (African strain), were treated with a range of dilutions of VIROSAL for 2 minutes and the viral load was quantified by qRT-PCR. (C) Zika virus (PE243) was treated with VIROSAL as above and used to infect human foreskin cells for 48h. Cells were stained with 4G2 anti-flavivirus antibody or an isotype control and anti-mouse Alexa-488 secondary antibody. Infected cells were quantified by flow cytometry. Statistical differences are presented relative to the pH5.5 control (n = 3 independent experiments). \*\**P* < 0.01, \**P*<0.05.

To investigate whether ViroSAL inhibits a broad range of enveloped viruses, viruses rescued from infectious clones of HSV-1, Measles, Epstein-Barr virus, and orf, a sheep parapox virus, were treated with ViroSAL at various concentrations and used to infect Vero cells, Vero cells engineered to express the measles entry receptor, SLAM, primary human tonsil epithelial cells and primary lamb skin epithelial cells, respectively. ViroSAL effectively inhibited infection of all viruses tested (Figure 3), and similar to Zika virus, a higher concentration of ViroSAL (3%) was required for complete neutralization of full-length viruses compared to pseudoviruses. Treatment with 3% ViroSAL did not result in cytotoxicity, measured using MTT assays (data not shown). This indicates that ViroSAL is capable of fully neutralizing a range of enveloped viruses *in vitro*, at a concentration that does not cause cytotoxicity in eukaryotic cells.

**Figure 3:**
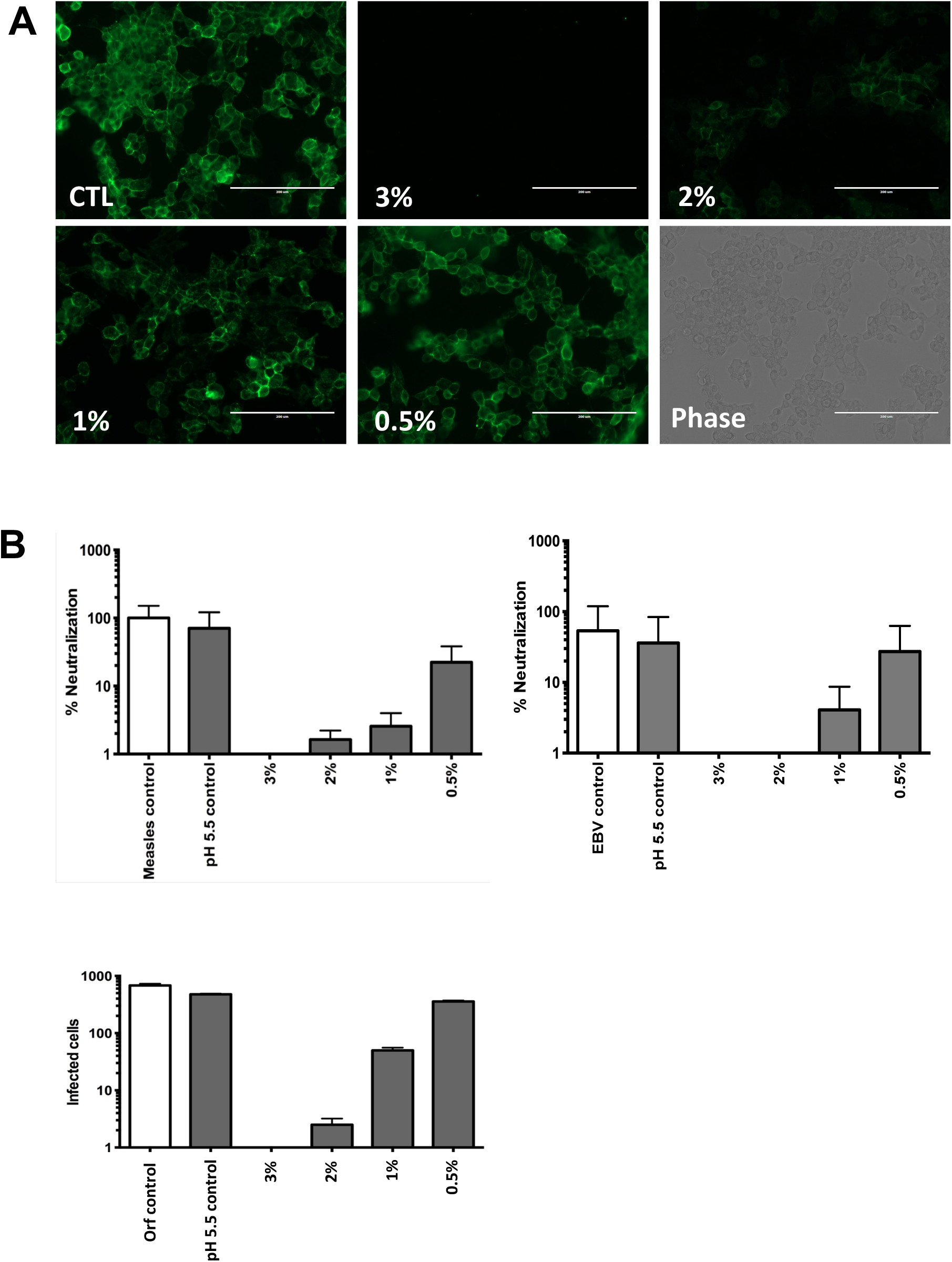
VIROSAL inhibits HSV-1, measles, Epstein-Barr and orf parapoxvirus infection. HSV-1 (strain I7) was treated with ViroSAL for 2 minutes, the pH restored to 7, and used to infect Vero cells. Control virus was treated with a pH5.5 buffer for 2 minutes and the pH restored to 7. After 24 hours, cells were fixed and stained with an antibody to detect HSV gD glycoprotein. Images are shown with a representative phase image. (B) Measles, EBV or orf virus was treated with VIROSAL for 2 minutes, the pH restored to 7, and used to infect Vero, primary tonsil epithelial cells or primary foetal lamb skin cells, respectively. Control virus was treated with a pH5.5 buffer for 2 minutes and the pH restored to 7. After 48 hours, viral infectivity was enumerated. Data are presented as % neutralization of infection relative to the untreated virus control.

### 1.3.4 ViroSAL does not inhibit infection of norovirus, a non-enveloped virus

Since ViroSAL potently inhibits infection of a range of enveloped viruses, we sought to establish whether this effect was limited to enveloped viruses or whether ViroSAL also influenced non-enveloped viruses. Murine norovirus, a non-enveloped virus, was treated with ViroSAL and added to cultures of RAW264.7 murine macrophages or BV2 murine microglial cells. There was no significant difference between control treated virus and ViroSAL treated virus infectivity in either cell type tested (Figure 4A). This indicates that, in contrast to ViroSAL’s effect on enveloped viruses, ViroSAL does not inhibit infection with norovirus, a non-enveloped virus.

**Figure 4:**
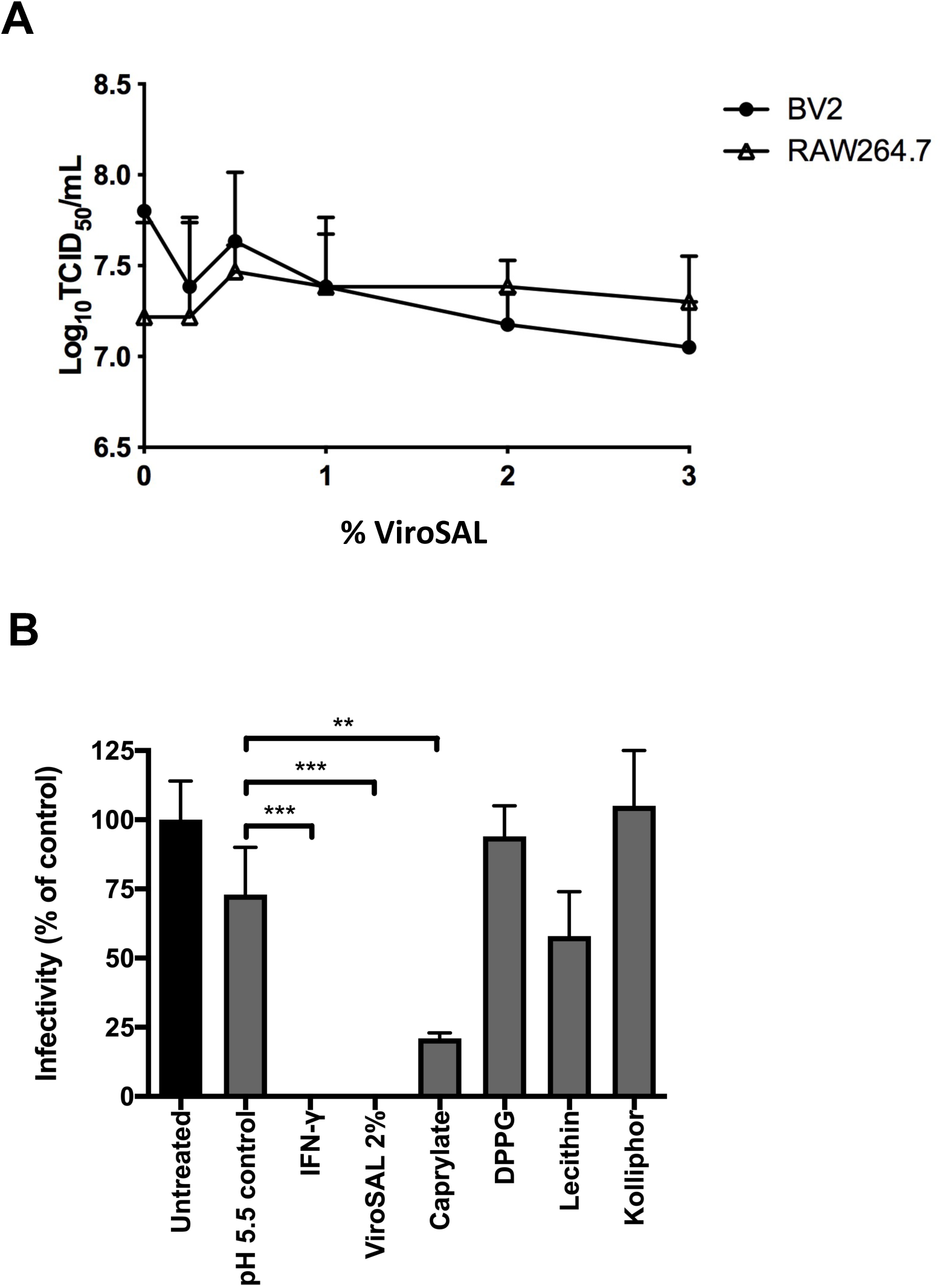
ViroSAL has no effect on the infectivity of non-enveloped norovirus, and caprylate is the major component responsible for the antiviral effect of ViroSAL. (A) Murine norovirus was treated with ViroSAL for 2 minutes, the pH restored to 7, and used to infect either RAW264.7 murine macrophages or BV2 murine microglial cells. After 96h, viral infectivity was measured and quantified as TCID50 via microscopic visualization. No significant difference was observed following ViroSAL treatment compared with control treated virus. (B) VSV-G pseudovirus was treated with 2% ViroSAL or the equivalent concentrations of caprylate, DPPG, Lecithin and Kolliphor for 2 minutes, and then pH restored to 7. To control for the effect of pH on viral infectivity, virus was treated with a pH5.5 buffer, equivalent to that of ViroSAL, for 2 minutes and then the pH of the virus was restored to 7. 10ng/ml IFN-y was included as a positive antiviral control. Data are presented as mean infectivity ± SD relative to the untreated virus control (n = 3 independent experiments). \*\*\**P* < 0.001, \*\**P* < 0.01, ns: not significant.

### 1.3.5 Caprylic acid is the major component of ViroSAL responsible for antiviral activity

To investigate the components of ViroSAL that are responsible for its activity, we evaluated individual components of ViroSAL (Caprylate, DPPG, Lecithin and Kolliphor), at concentrations equivalent to amounts present in 2% ViroSAL, together with 2% ViroSAL and 10ng/ml IFN-y as a positive antiviral control. Only Caprylate (caprylic acid) had a significant effect on VSV-G pseudovirus entry to Vero cells (Figure 4B). However, 2% ViroSAL fully neutralized viral entry, indicating that caprylic acid within the ViroSAL formulation was more potent as an antiviral than caprylic acid alone.

### 1.3.6 ViroSAL disrupts orf virion integrity

To examine the potential perturbation of enveloped virus membranes, we used transmission electron microscopy to visualize ViroSAL-treated orf virions. Orf is a parapoxvirus, a family of viruses that includes pseudocowpox and bovine papular stomatitis virus, and it was selected because of the large size of the virions (200-300nm in diameter) and the relatively high titres that can be achieved following growth in cell culture. Negatively stained virions exhibited typical poxvirus morphology (Figure 5A), with an ovoid structure surrounded by an external envelope. Treatment of orf virions with 1% ViroSAL, which neutralized approximately 50% infection in infection assays (Figure 3B), appeared to disrupt the external viral envelope, which remained associated with the virions (Figure 5B). Treatment with 3% ViroSAL, which completely neutralized *in vitro* infectivity (Figure 3B) disrupted virion morphology and the lattice-like structure present on intact virions (Figure 5C). These data indicate that ViroSAL disrupts orf surface structure, and virion morphology.

**Figure 5:**
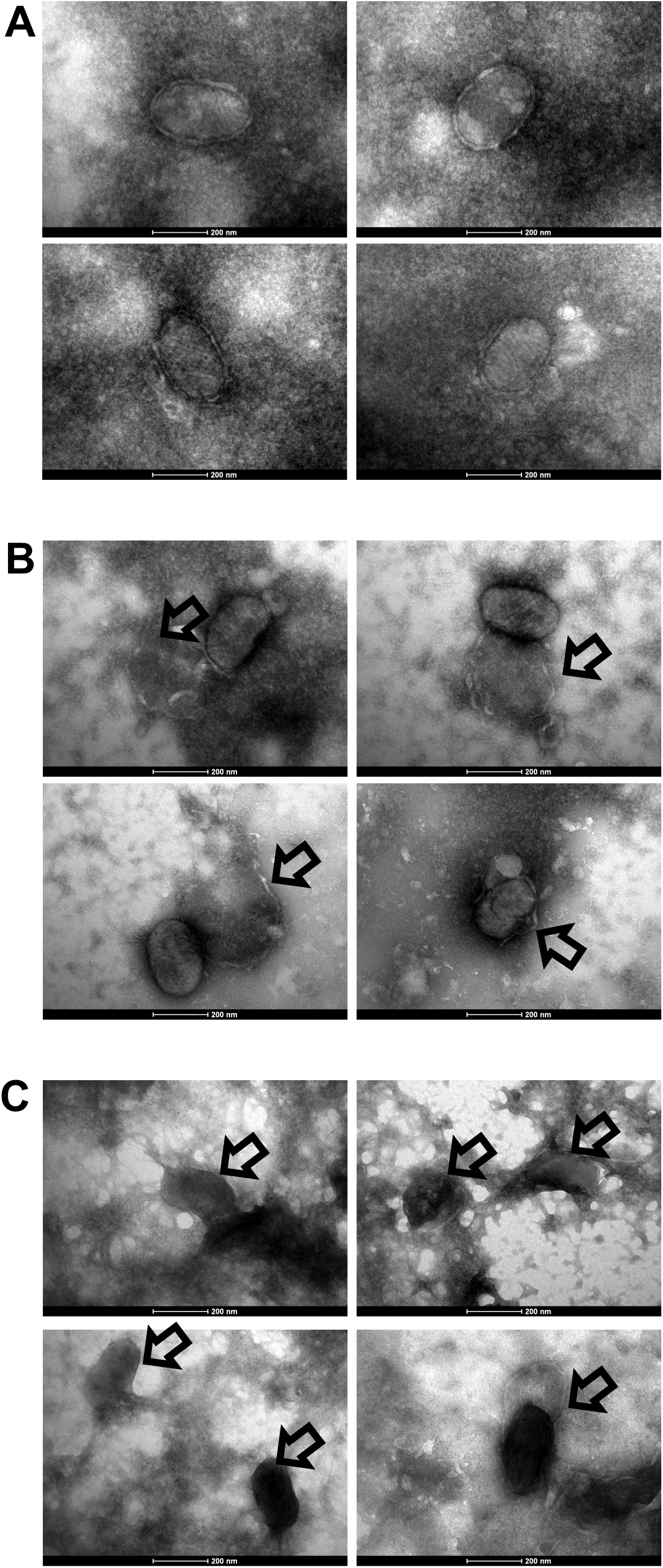
ViroSAL treatment of Orf virus disrupts the structural integrity of virions. Orf virus (MRI-scab) was treated with a pH5.5 buffer (A), 0.5% ViroSAL (B) or 3% ViroSAL (C) for 2 minutes, and pH was restored to 7. Mock treated virions exhibited typical poxvirus morphology with a prominent outer envelope (A). Virions treated with a low concentration of ViroSAL (1%) displayed altered morphology and external envelope integrity was disrupted (arrows)(B). Virions treated with a high concentration of ViroSAL (3%) exhibited heterogeneous morphology with extensive disruption of virion structure (arrows).

### 1.3.7. ViroSAL inhibits Semliki Forest virus and Zika virus *in vivo*

Arboviruses are transmitted to their mammalian hosts via the bite of an insect vector. Given that ViroSAL has optimal activity at a slightly acidic pH, such as that seen on skin and mucosal surfaces, we investigated the effects of ViroSAL treatment on mouse skin following viral inoculation at the site of *Aedes aegypti* bites, according to previously published protocols (Bryden et al., 2020). Arbovirus replication in the skin at mosquito bites is a key stage of infection during virus replicates rapidly before disseminating to the blood and other tissues (Pingen et al., 2016). Previous work has suggested therapeutic targeting this site may be efficacious in reducing severity of infection (Bryden et al., 2020). Here, ZIKV was chosen, as it is a medically important emerging arbovirus, whereas SFV was used as a model virus that replicates efficiently in immunocompetent mice.

To firstly show that ViroSAL can inactivate infectious SFV *in vitro* (Supplementary Figure 2) in addition to ZIKV (Figure 2), SFV6 was incubated at increasing infectious titres with 5% ViroSAL at pH5.5 for 2 minutes. Following immediate restoration of physiological pH, solution was applied to monolayers of BHK-21 cells and infectious titre assessed by plaque assay. As a control SFV6 was incubated similarly at pH5.5 in the absence of ViroSAL. The amount of infectious SFV was reduced to beyond the limit of detection by ViroSAL, suggesting this virus is highly sensitive to treatment (Supplementary Figure 2). Similarly, when ViroSAL treated SFV6 was inoculated subcutaneously into mice, the titre of virus at 24 hours post infection was significantly reduced (Supplementary Figure 2).

We next determined whether topical application of ViroSAL can also suppress virus infection when virus was inoculated subcutaneously into a mosquito bite. In this experiment, ViroSAL was applied immediate post mosquito bite and left for 20 minutes to allow penetration of skin by ViroSAL, or left untreated as a control. SFV was then inoculated subcutaneously into the mosquito bite as previously described (Pingen et al., 2016). Re-application of ViroSAL to the inoculation site was repeated at 5 hours post infection. At 24 hours post infection, those mice receiving ViroSAL had a significantly reduced amount of virus RNA at the inoculation/bite site and limited dissemination of virus to the spleen 24 hours post infection (Figure 6A). Infectious viral titres within plasma were also significantly reduced (Figure 6B). Topical application of ViroSAL to mosquito bites was similarly efficacious in reducing ZIKV serum viraemia by 24 hours post ZIKV infection (Figure 6C).

**Figure 6:**
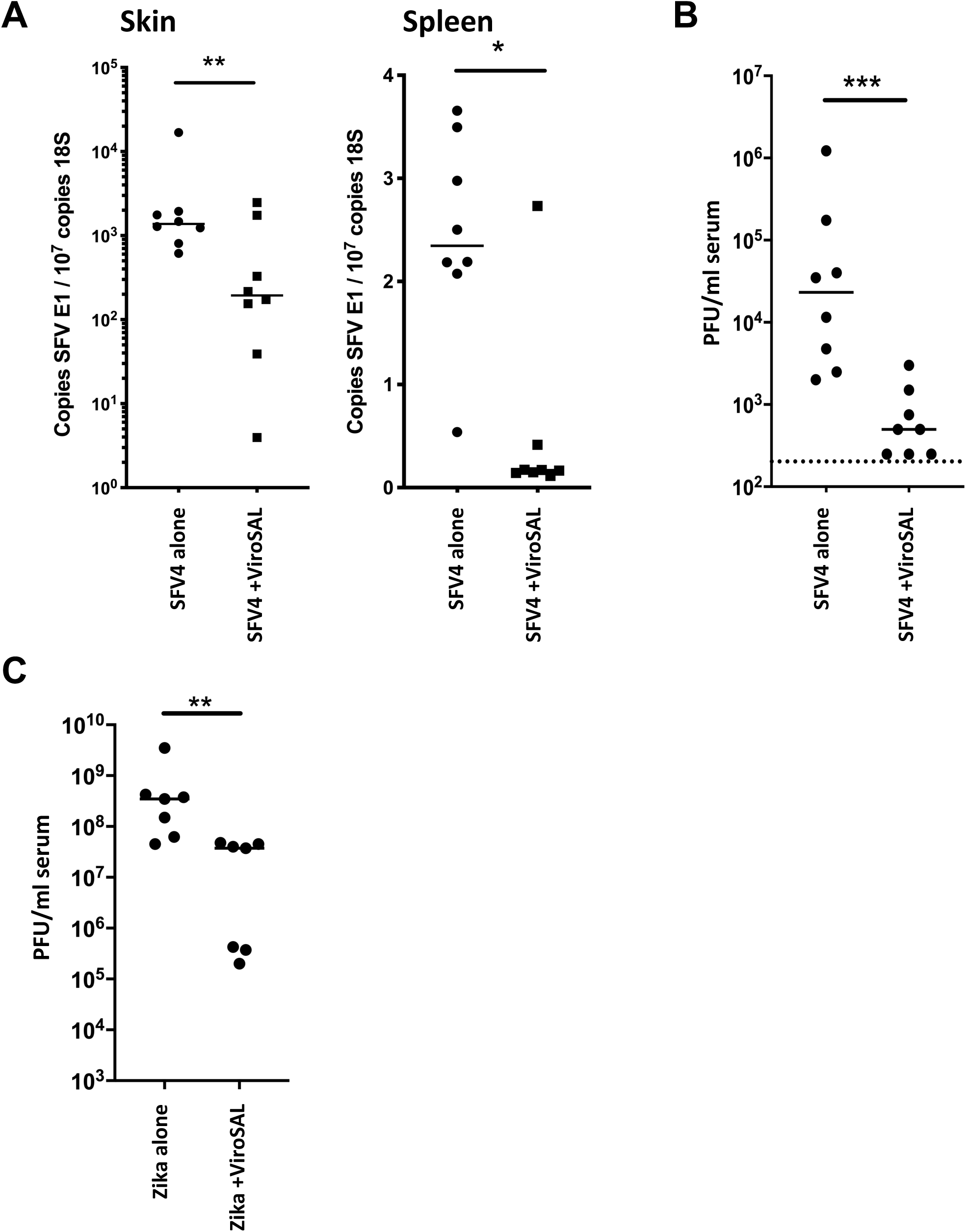
ViroSAL inhibits local and systemic replication of SFV and ZIKV *in vivo*. (A) Following a bite by an *Aedes aegypti* mosquito of a mouse foot (upper side), skin was treated with topical ViroSAL. Following 20 minutes, to allow for penetration of skin by ViroSAL, the same site was inoculated with Semliki Forest virus (SFV4) or Zika virus by microneedle injection. The bite site was re-treated with ViroSAL at 5 hours post inoculation (8 mice) or left untreated (8 mice). 24 hours post infection, animals were euthanized and viral loads in skin and spleen enumerated by qRT-PCR. (B,C) Infectious virus from SFV4 and Zika infected mice was enumerated by plaque forming assay (PFU) 24 hours post-infection. *P < 0.05, **P < 0.01, ***P < 0.001 (Mann-Whitney test).

## 1.4 Discussion

The ViroSAL used in these studies is a proprietary emulsion of free fatty acids that has previously been shown to have antibacterial properties, including against a range of veterinary periodontopathogens (Laverty et al., 2015), *Staphylococcus* species including methicillin-resistant *Staphylococcus aureus* (Hogan et al., 2016) as well as a wide range of culturable species from human colon (McDermott et al., 2015). The antimicrobial activity of medium chain saturated and long chain unsaturated free fatty acids has been reported previously (Thormar and Hilmarsson, 2007), with both antibacterial and antiviral activity observed (Hilmarsson et al., 2005).

Caprylic acid has been used in the purification process of immunoglobulins due to its inactivation of lipid enveloped viruses (Dichtelmuller et al., 2002), a process which is pH-dependent and optimal at low pH. Poor solubility in aqueous solutions has limited antimicrobial applications of fatty acids in their free form (Jackman et al., 2016) and dissolution in solvents such as DMSO can cause skin irritation. Formulation as nanoparticles has been described to overcome the challenges of antimicrobial fatty acid bioavailability (Jackman et al., 2016, Yoon et al., 2018). In this study we used patented technology (Folan, M.A. International Patent No. WO 2011/061237) to build and use an emulsion to evaluate its antiviral potential. Recent publications have noted the potential antimicrobial activity of nanoemulsions and microemulsions (Churchward et al., 2018, Buranasuksombat et al., 2011, Ma et al., 2016, Donsi and Ferrari, 2016). In each of these studies there are no data that clearly evaluate the contribution of the individual components to the overall properties which, in a mixed system may be synergistic or antagonistic. However, each of the components of ViroSAL is Generally Regarded As Safe (GRAS) by the United States Food & Drug Administration.

The antiviral activity of ViroSAL emulsion may be due to amplified surface area of the water insoluble fatty acid together with the amphipathic nature of the de-lipidised lecithin which facilitates delivery to lipophilic cell surfaces and/or the viral envelope (Laverty et al., 2015). We observed a significant concentration- and time-dependent (rapid) decrease in infectivity of all enveloped viruses tested in the present study, with no effect on non-enveloped norovirus. Milk-based free fatty acids, as well as fatty acid emulsions, have been shown to inhibit infection of Vero cells with VSV and HSV-1, with no antiviral effect on poliovirus, a non-enveloped virus (Thormar et al., 1987). Moreover, Isaacs (Isaacs et al., 1990) reported no antiviral activity of human milk or milk formula, but potent antiviral activity of milk aspirated from the stomachs of infants one hour after feeding, which occurred due to the release of free fatty acids by lipolysis. Similar observations were reported by Thormar et al. (Thormar et al., 1994), after lipolysis by storage of human milk at 4°C. Following treatment of orf virus with ViroSAL, we observed a dissociation of the viral envelope from virions, with higher concentrations disrupting virion integrity and morphology. In agreement with the present study, fatty acids were found to affect the viral envelope, with high concentrations causing complete disintegration of viral particles (Thormar et al., 1987, Thormar et al., 1994). In agreement with the present study, previous studies have also reported no antiviral effect of fatty acids on non-enveloped viruses (Thormar et al., 1987, Churchward et al., 2018), indicating that the antiviral mechanism of free fatty acids and ViroSAL may involve the viral envelope and we anticipate that to be due to a surfactant effect (Yoon et al., 2015, Yoon et al., 2017).

The non-ionised form of caprylic acid dissociates at basic pH into the ionized form and only the non-ionised form is capable of virus inactivation (Lundblad and Seng, 1991). ViroSAL has optimal activity at pH5.5 and is consequently suitable for use in environments close to this pH, such as skin, oral cavity and mucous membranes which are also relevant portals of infection. The present study demonstrated that ViroSAL has antiviral activity against HSV-1, EBV and orf viruses, all of which are pathogens of skin or mucous membranes and have a tropism for epithelial cells. Since ViroSAL has optimal activity at pH5.5, we also tested a range of epithelial cell systems that would physiologically have a lower pH than that of plasma, such as skin which has a pH of approximately 5 (Lambers et al., 2006) and the oral cavity which can have a pH of 6.5 or lower depending on the level of oral health. ViroSAL had strong antiviral activity against Semliki Forest and Zika viruses, which are transmitted via mosquitoes into the skin dermis. This is an important stage of the virus life cycle, in which it replicates in dermal cells before disseminating to the blood and then throughout the body (Hamel et al., 2015; Pingen et al., 2017). The potential of ViroSAL as a topical application immediately following mosquito bites is of great interest, as established and emerging mosquito-borne viruses constitute an increasing threat to human health. Treatment modalities that incorporate ViroSAL to target skin arbovirus infection should be explored further as e.g. treatment of the mosquito bite inoculation site with an immunomodulator has recently been shown to be an effective strategy in suppressing arbovirus infection (Bryden et al., 2020). Moreover, ViroSAL also neutralized measles virus and SARS-CoV-1 pseudovirus infection. Measles is an aerosol transmitted infection that initially targets macrophages and dendritic cells of the upper airway, which are then transported across the respiratory epithelium to the lymphatic organs (Muhlebach et al., 2011). Similarly, SARS-CoV-1 causes an acute interstitial pneumonia and was the causative agent of the 2002 SARS coronavirus epidemic. ViroSAL, therefore, has potential applications as a topical or inhaled therapy to treat epitheliotropic viral infections or those that infect the airway or other mucous membranes. The fact that ViroSAL has applications against a wide range of enveloped viruses is of great interest, and while many antivirals have activity against specific viral families, there are few broad-spectrum antivirals currently available. Therefore, these data provide a rationale for future pre-clinical studies to investigate the antiviral properties of ViroSAL in viral infections of mucous membranes including the airway and skin. ViroSAL may have applications in current and future viral outbreaks, including the current pandemic caused by SARS-CoV-2.

## Acknowledgements

We acknowledge Dr. Colin Crump, University of Cambridge, for providing HSV reagents, Prof. Alain Kohl, University of Glasgow, for providing Zika reagents and Dr Colin McInnes and Lesley Coulter at the Moredun Research Institute, Midlothian, Scotland, United Kingdom for providing orf virus and foetal lamb skin cells. We also thank Sandra Terry and Dr Emilie Pondeville for supplying *A. aegypti* mosquito eggs; and the University of Leeds St James’ Biomedical Services and Gillian Cardwell for assistance with mouse studies.

