## Supplementary figures. for "A novel antiviral formulation inhibits a range of enveloped viruses"

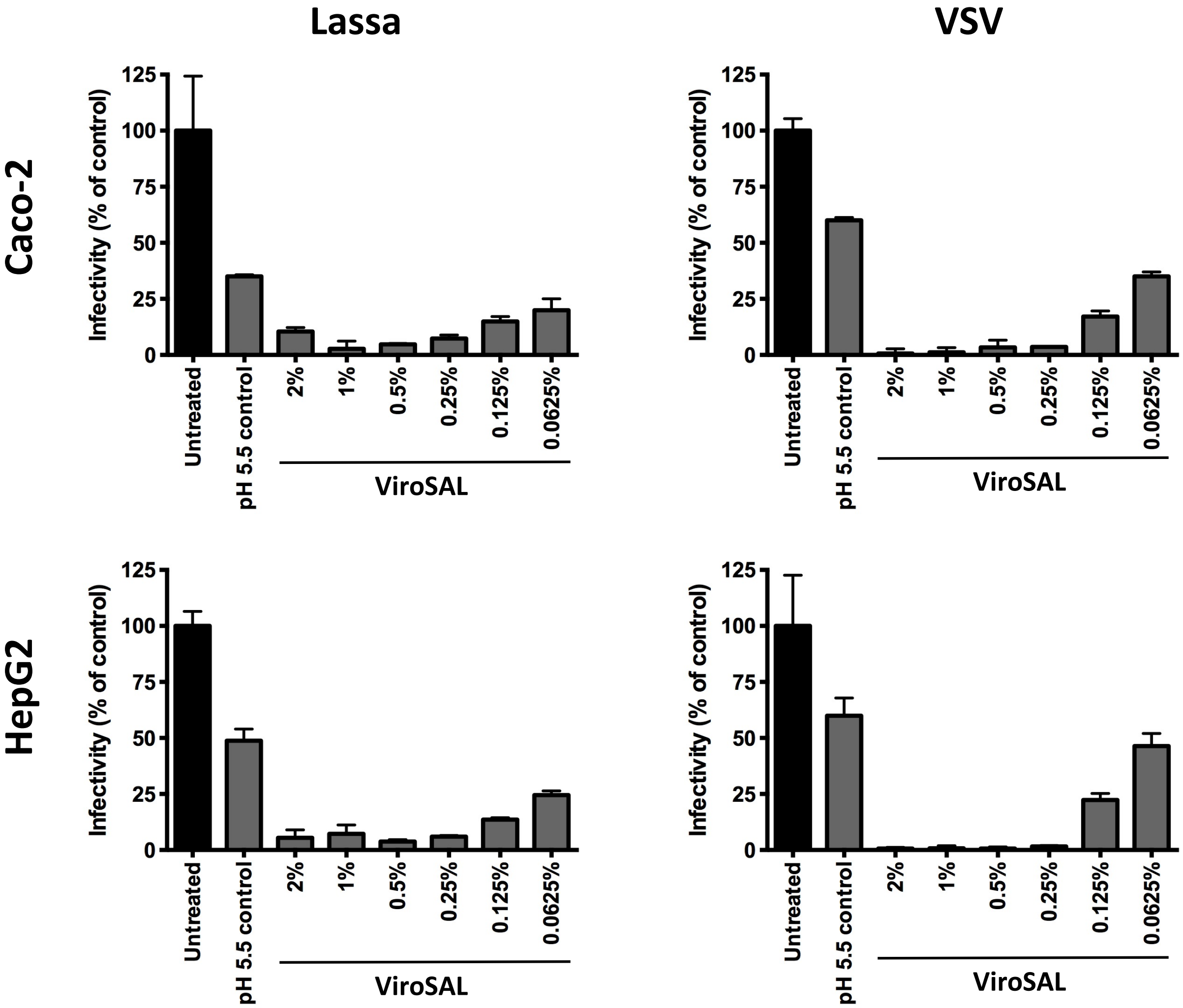


**Supplementary Figure 1: ViroSAL inhibits cellular entry of pseudoviruses in multiple cell lines.** Pseudovirus bearing the envelope glycoproteins of Lassa or VSV was treated in a 1:1 dilution with concentrations of ViroSAL ranging from 4% to 0.125% (final concentrations ranged from 2% to 0.0625%) for 2 minutes. Buffer was then added to restore the pH to 7. To control for the effect of pH on viral infectivity, virus was treated with a pH5.5 buffer, equivalent to that of ViroSAL, for 2 minutes and then the pH of the virus was restored to 7. Pseudoviruses were used to infect 293T human embryonic kidney cells. Data are presented as mean infectivity ± SD relative to the untreated virus control.

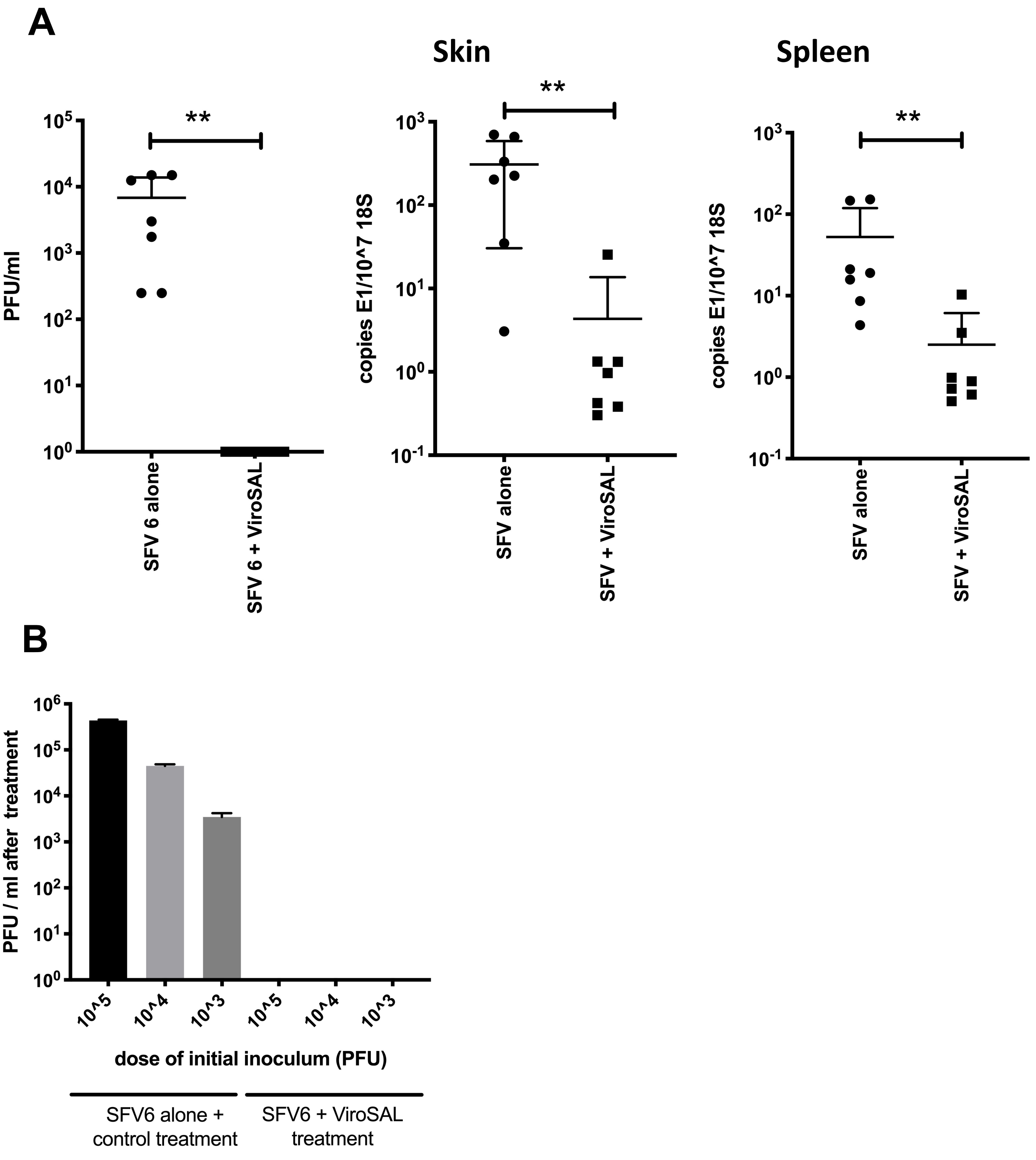


**Supplementary Figure 2. ViroSAL inhibits SFV6 *in vitro and in vivo*.** (A) **Semliki Forest virus (SFV6)** was treated in a 1:1 dilution with 10% ViroSAL (final concentration 5%) for 2 minutes. Buffer was then added to restore the pH to 7. Virus was added to Vero cells and infection enumerated by measurement of plaque forming units (PFU; first panel) or injected subcutaneously into wild type C57/bl6 mice, and viral loads in skin and spleen quantified by qRT-PCR. (B) SFV6 was treated with a 1:1 dilution 10% ViroSAL (final concentration 5%) for 2 minutes. Buffer was then added to restore the pH to 7. Virus was added to Vero cells and infection enumerated by measurement of plaque forming units (PFU). **P < 0.01 (Mann-Whitney test).
